# Retention of the full visual opsin repertoire in Australia’s cryptozoic lizards

**DOI:** 10.1101/2024.02.19.581082

**Authors:** Matthew J.R. Ford, Alastair J. Ludington, Tessa Bradford, Kate L. Sanders, Mark N. Hutchinson, Bruno F. Simões

**Author notes:** Email addresses: Alastair Ludington Tessa Bradford Kate Sanders Mark Hutchinson. Corresponding Authors: Matthew J.R. Ford, Bruno F. Simões.

## Abstract

Australian scincid lizards in the sister-genera *Lerista* and *Ctenotus* are a prominent system for understanding adaptation in the transition from surface to fossorial life. The approximately 205 species in this group exhibit extreme diversity in morphology and ecology. *Lerista* and *Ctenotus* both include diurnal and surface-active species that are fully pentadactyl, and *Lerista* also contains many specialised limb-reduced and limbless sand-swimmers. To understand how the visual systems of these lizards have responded to their varied photic environments, we examined the five opsin genes encoding the pigments that mediate colour and dim-light vision. These genes were sequenced for 59 species of *Lerista* and *Ctenotus* and analysed for variation in selection pressures among amino acid sites and across branches in the species tree. All five opsins are present and intact in all species of *Lerista* and *Ctenotus* examined, and we identified signals of positively selected substitutions in all five opsin genes –RH1, which mediates scotopic vision, and four cone opsins associated with photopic vision (SWS1, SWS2, RH2, LWS). Most comparisons of selection pressures did not show significant differences according to broad ecological divisions. Only LWS showed a signal of relaxed selection in sand-swimming (limb reduced) versus less fossorial (fully limbed) *Lerista*. These results suggest that photopic abilities are retained across both clades, even in the most fossorial species, highlighting a need for studies of visual ecology of Australian skinks, and prompts caution with regards to generalisations about degenerate vision in fossorial squamates.

## Introduction

Evolutionary transitions from bright-light (photopic) to low-light (scotopic) environments are important drivers of change in the evolution of visual systems (Walls, 1942; Zhao et al, 2009; Pinto et al, 2019). In many taxa that rely on vision for vital tasks such as feeding, predator avoidance and mating, reduced light availability is compensated by increased corneal diameter and visual sensitivity (Hall and Ross, 2007; Hall, 2008). However, while indispensable for many taxa, eyes and optic neural tissue are also energetically expensive to develop and maintain, and these costs are often linked to evolutionary degeneration of eyes and vision in scotopic habitats (Moran et al, 2015; Protas et al, 2007). Indeed, the repeated loss of visual function in scotopic-adapted mammals, reptiles, amphibians, fish and insects is a prominent example of convergent evolution (Sadier et al, 2018; Mohun et al, 2010; Simões and Gower, 2017; Musilova et al, 2021; Sharkey et al, 2017). In these taxa, the degree of visual conservation or degeneration is expected to vary with ecological context. The subcutaneous eye of the obligate fossorial *Spalax* mole-rats is retained only for a non-visual, circadian function (Avivi et al 2002; Haim et al, 1983), whereas the primarily subterranean blind snakes (Simões et al, 2015; Gower et al, 2021) and star-nosed moles (Emerling and Springer, 2014) have degenerated eyes and reduced colour discrimination.

Transitions to new photic environments can also be accompanied by changes to the visual pigments that activate the phototransduction cascade (Cronin et al, 2014) and are encoded by opsin genes (Emerling and Springer; 2014; Simões et al, 2015; 2020; Hauser et al, 2021). In the ancestral tetrapod (and squamate) visual system, five spectrally distinct opsin pigments are expressed in two main photoreceptor types (Yokoyama, 2000; Solomon and Lennie, 2007). Cone photoreceptors are primarily responsible for photopic vision and contain short (∼360nm) to long (∼600nm) wavelength sensitive opsins that enable colour discrimination; rod photoreceptors contain rhodopsin 1 (RH1) and are used for vision in scotopic conditions (Walls, 1942; Crescitelli, 1977; Yokoyama, 2000; Bowmaker, 2008; Hunt et al, 2009; Davies, 2011). Transitions to scotopic environments are thought to be linked to selective losses of cone opsins (and associated phototransduction genes), and preferential retention of rod-expressed opsins, in several lineages of snakes, crocodilians, mammals and amphibians (Simões et al, 2015; Gower et al, 2021; Emerling, 2017; Emerling and Springer, 2014; Mohun and Davies, 2019). In squamates, this has occurred in geckos (Pinto et al, 2019) and snakes (Simões et al, 2015, 2016; Gower et al, 2021). Gene losses occur when purifying selection is reduced due to a relaxation of selective constraints, resulting in the accumulation of inactivating mutations. In some echolocating bats, evidence of relaxed selection on the SWS1 opsin is associated with the loss, or ‘pseudogenisation’, of this gene (Wertheim et al, 2015; Simões et al, 2019). Alternatively, cone opsins might be intact but evolve at accelerated rates indicative of relaxed selective constraints in low-light environments (Emerling and Springer, 2014; 2015). Where opsins are retained and their function is conserved, purifying selection is expected to remove harmful variants (Veilleux et al, 2013). Alternatively, positive selection might favour an adaptive variant that changes some aspect of opsin function, such as wavelength of maximum absorbance. These signals of positive selection are often detectable by the amino-acid complement at so-called spectral tuning sites that influence spectral sensitivity (Fasick & Robinson, 1998; Cowing et al, 2002; Yokoyama, 2008a and citations therein).

Clades of closely related taxa that have undergone parallel transitions between photic environments provide powerful systems for understanding visual evolution during ecological transitions. Australian scincid lizards in the genera *Lerista* and *Ctenotus* are an attractive system in this respect. The ∼205 species (Uetz et al, 2022) in this clade share a common ancestor ∼31 million years ago (Rabosky et al, 2014; Skinner et al, 2013; Hedges et al, 2015; Zheng et al, 2016; Tonini et al, 2016; Pyron et al, 2014; Wiens et al, 2006; Wright et al, 2015). All *Ctenotus* are surface active, with unreduced, pentadactyle limbs, little tendency to body elongation, and well-developed eyes with moveable scaly lids (Greer 1989). While all species are primarily day-active in bright light desert environments, some, most notably *Ctenotus pantherinus*, show increased crepuscular and (less commonly) nocturnal activity or occupy light-limited habitats such as leaf litter in mesic woodlands (Gordon et al, 2010). In contrast to *Ctenotus*, *Lerista* species are highly diverse in habitat and morphology (Cogger, 2014). Across *Lerista*, there have been many independent adaptations of morphology to better suit a more cryptozoic lifestyle, such as elongation of the body (Mann, 2020), limb reduction and digit loss (Greer, 1987; Greer, 1990; Skinner et al, 2008), and alteration of the skull (Pough et al, 1997) (see also Fig. 2). Some lineages are more diurnal and surface-active, and although showing some loss of digits, they retain four functional limbs, whereas several independent lineages are more fossorial and show reduction in relative eye size and more extreme burrowing (“sand-swimming”) adaptations such as loss or extreme reduction of forelimbs, loss of all digits with forelimbs absent and hind limbs styliform or absent, and further increase in vertebral number (Storr, 1976; Storr, 1990; Greer, 1987; Greer, 1990; Skinner et al, 2008; Couper et al, 2016). While the ancestral state of *Lerista* is not definitively known, it is thought that the ancestor of the genus was tetrapodal and probably pentadactyl (Morinaga and Bergmann, 2017), as re-acqusition of a fully pentadactyle limb from a greatly reduced state is unlikely (Skinner et al, 2008; Skinner, 2010; but see also Wagner et al, 2018; Bergmann et al, 2020).

The repeated shifts in photic environments during the evolution of *Lerista* provide a valuable opportunity to identify adaptive responses of visual systems to environmental factors. We might hypothesise that the opsin genes of highly fossorial skinks will show signals of relaxed selection or regressive evolution. However, we emphasise that these skinks must experience complex and variable selection pressures on their visual systems, given the observation that few are known to have identical combinations of limb morphology, eye-lid condition, activity patterns and habitat use (Fig. 1).

**Figure 1:**
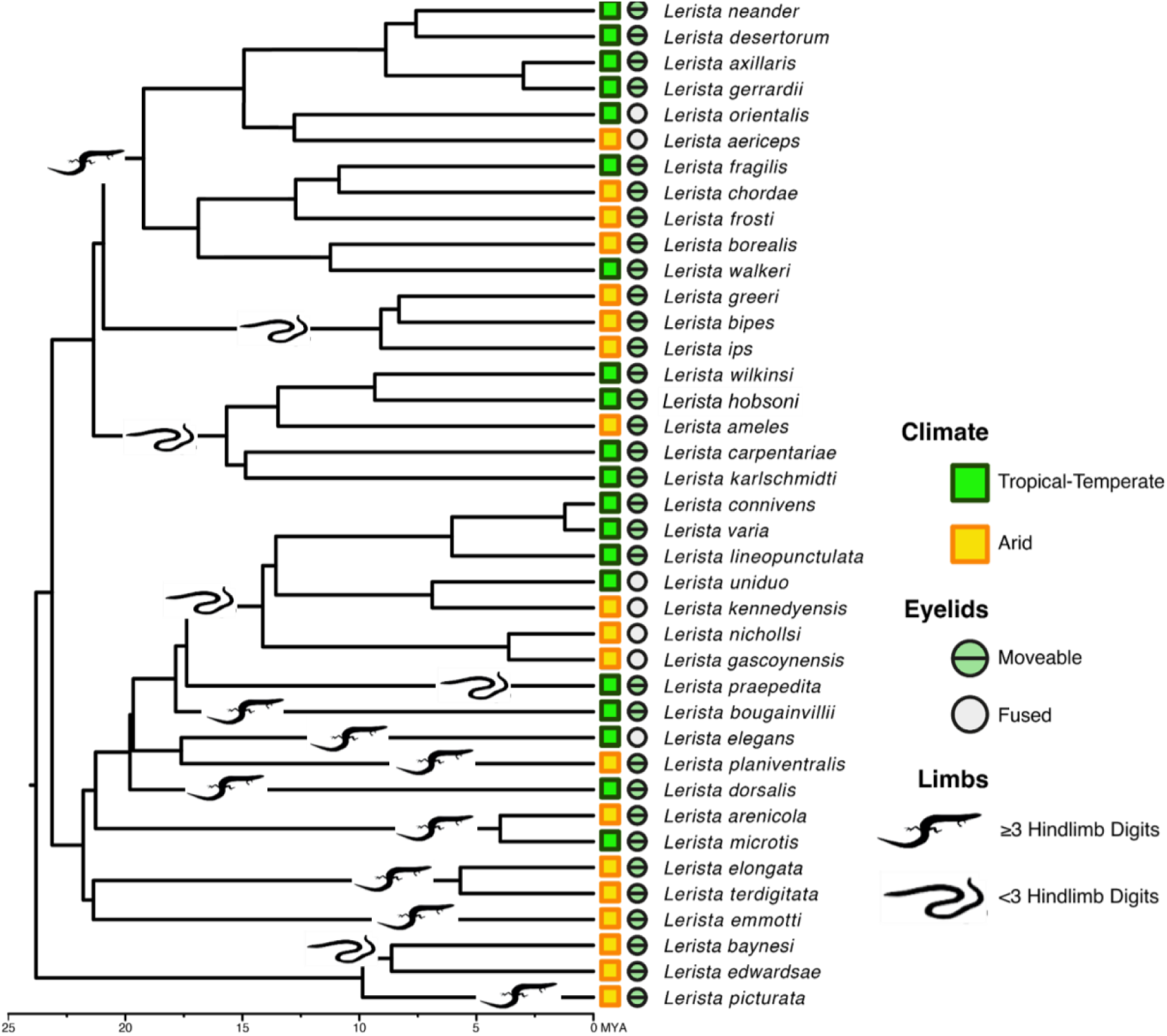
Phylogenetic tree of *Lerista* showing divergence time estimates (in MYA) of the species included in this study and their habitat and limb state. Time tree obtained from VertLife phylogeny subsets.

**Figure 2:**
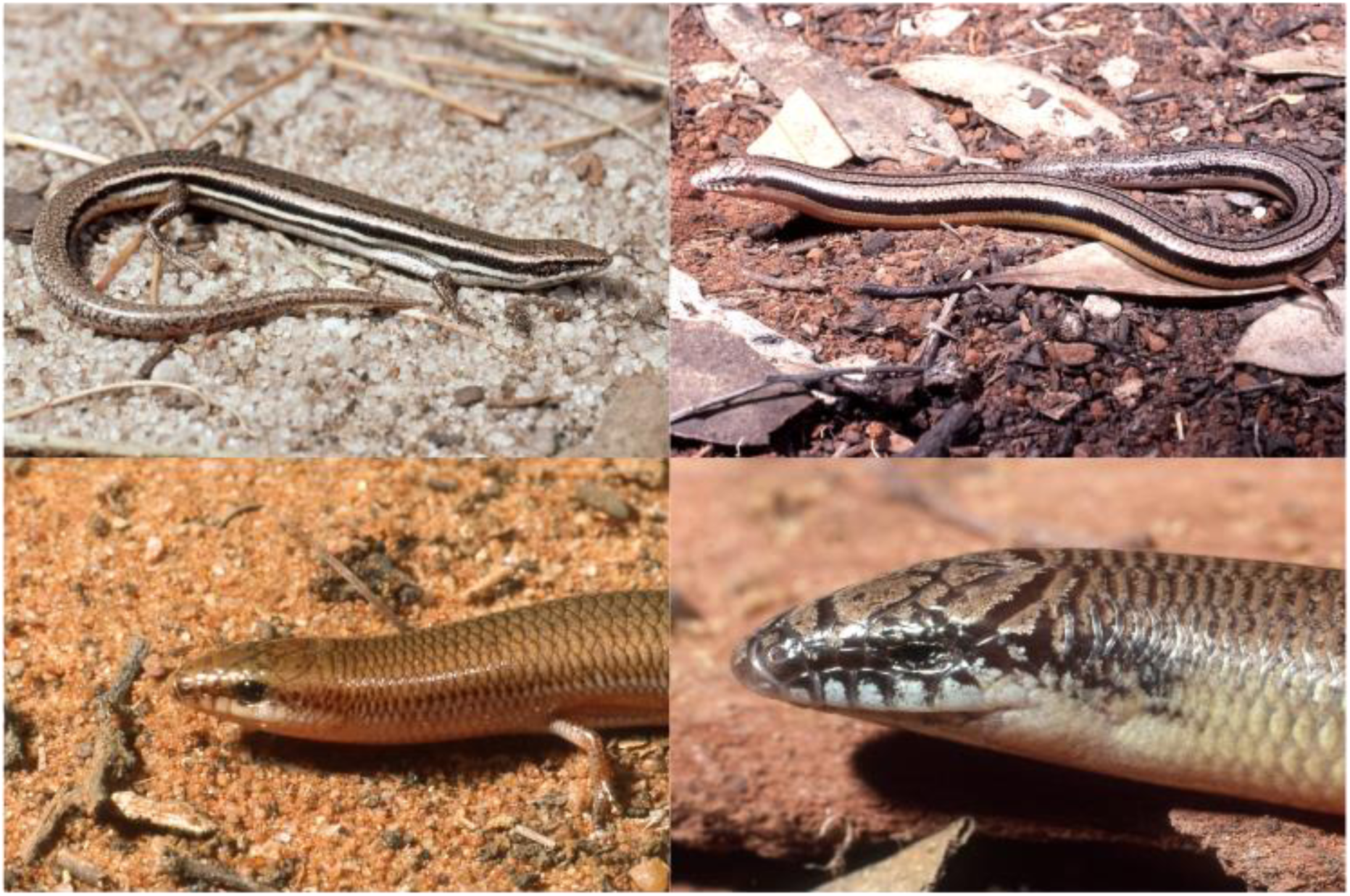
Four *Lerista* showing different limb and eyelid states. Clockwise from top left: *Lerista microtis*, tetrapodal and pentadactyl; *Lerista edwardsae*, hind limbs only, bidactyl; *Lerista punctatovittata*, showing small eyes with movable eyelids; *Lerista aericeps*, showing large eyes with immovable eyelids modified as a spectacle covering the eye. © Mark Hutchinson

This study used molecular analyses to explore the diversity of visual traits in *Lerista* and *Ctenotus* and, more specifically, investigate opsin evolution during transitions to fossorial habits in *Lerista.* In these respects our paper provides a substantial contribution to the literature on the nature of visual opsin adaptations to low light conditions in Squamata. We are among the first to analyse the visual opsin complement of Scincidae, a previously understudied squamatan clade with regards to visual pigment evolution.

## Methods

### Taxon sampling and DNA Extraction

We sequenced all visual opsins across 59 species of *Lerista* and *Ctenotus*, including diverse surface-active species and spanning multiple lineages that have independently acquired transitional or highly fossorial habits (Skinner et al, 2008, Fig. 1). This sampling allowed us to perform comparative tests of opsin selection across repeated transitions in habitat and diel activity. The recent timing of these transitions, estimated from 3.6 to 11.8 million years (Skinner et al, 2008), reduces the possibility of diversity in opsins being due to factors not causally associated with the transition in photic environment (Rabosky et al, 2007; Skinner et al, 2013; Rabosky et al, 2014). In total,we extracted DNA for 127 species in 23 genera of skinks (Squamata: Scincidae) obtained via fieldwork (35 samples) and from The Australian Biological Tissue Collection (ABTC) of the South Australia Museum (92 samples) (SI Table 1). Additionally, we extracted total RNA from the eyes of 13 species for transcriptome sequencing (SI Table 1). Fieldwork and tissue collection was approved by permits from the Animal Ethics Committee of the University of Adelaide (21877) and from the Government of South Australia (Q26642-5) and Queensland (WA0009193). In addition, we downloaded the opsin sequences of other amniotes as outgroups from GenBank (SI Table 1).

DNA was extracted from liver and muscle using DNeasy Blood and Tissue kits (Qiagen, Chadstone, VIC, Australia) and following the manufacturer protocols. Samples were quantified using a QuBit^TM^ 2.0 (ThermoFisher Scientific, Waltham, MA, USA).

### Calculation of relative eye size

Relative eye size was calculated for most of the species using CT scans of specimens from the South Australian Museum, the Western Australian Museum and the Queensland Museum. We obtained CT scans using the Bruker Skyscan 1076 or 1276 in-vivo X-Ray microtomograph at Adelaide Microscopy, University of Adelaide. Specimens were scanned to provide volumes with voxel sizes of between 0.0085 and 0.017 mm depending on overall specimen size. All scans were made using a 0.5 mm aluminium filter to reduce scattering artefacts. The source voltage and current were both adjusted depending on the specimen size and ranged between 40-74 kV and 110-200 μA. Reconstruction and visualisation of slices was made using Bruker software (NRecon, DataViwer, CTAn) which produced image sets that are symmetrical in three dimensions and orthogonally oriented on the X,Y and Z axes. By viewing (CTAn, Bruker) the image stacks along the Z axis, the slices representing the first and last appearances of the skull and the scleral ring were noted, and converted into a measurement using the known thickness of the slices in each scan multiplied by the number of slices. Eye size (anteroposterior diameter of scleral ring) relative to skull length (mid line premaxilla-occipital condyle) values were log-transformed and plotted. Limb reduction was expressed using hind-limb digit counts as a proxy (SI Table 7). (could put a table into the supp)

### Library preparation and sequencing

Genomic DNA was sheared to ∼400-500bp using a Bioruptor Pico Sonicator (Diagenode, Denville, NJ, USA), and used a HS D1000 ScreenTape run on a Tapestation 2200 (Agilent Technologies, Santa Clara, CA, USA) to check the size distribution. Fragments over 200bp were selected for using Ampure XP (Beckman Coulter Inc, Brea, CA, USA) with 8.5% PEG for bead clean-up (Li et al, 2013). DNA was then prepared for Illumina sequencing based on the protocol of Meyer and Kircher (2010), using the on-bead method of Fisher et al (2011) and Li et al (2013). The DNA library underwent blunt end repair followed by adaptor ligation and ligation fill-in and the samples were double indexed by PCR amplification (Meyer and Kircher, 2010). The libraries were combined in equimolar amounts in pools of five samples each and then centrivapped down to 7µl and hybridised to custom RNA baits (Daicel Arbor Biosciences, Ann Arbor, Michigan, USA). We used MyBaits targeted capture protocols version v4 (http://www.arborbiosci.com/mybaits-manual/) with baits designed on the *Anolis*, *Pogona*, *Python* and *Gecko* genomes, where targeted vision genes were PCR-amplified in 16 cycles after capture with streptavidin beads (Dynabeads, ThermoFisher). Once amplified, these pools were all checked for concentration with the Qubit and fragment size distribution was measured again using a HS D1000 ScreenTape run on a Tapestation 2200 (Agilent Technologies, Santa Clara, CA, USA). The capture libraries were pooled in equimolar quantities to give a single sample at 10 nM DNA with an average fragment size of 400 bp. Illumina NovaSeq SP sequencing using 50bp paired end 100 cycle runs (single lane) were performed by Australian Genomics Research Facility (AGRF).

### RNA extraction, library preparation

Eleven RNA-seq samples were selected to provide reference sequences that are representative of the phylogenetic breath of the gene capture samples (SI Table 1). Eyes were collected via fieldwork and the eyes stored in RNAlater and frozen at -80°C (see SI Table 1). The eyes were macerated in TRIzol and the RNA was purified using the PureLink^TM^ RNA Mini Kit (ThermoFisher Scientific, Waltham, MA, USA) using the manufacturer’s protocol. The mRNA-Seq library was double-indexed and prepared with the mRNA-Seq Library Prep Kit v2 (Lexogen, Vienna, Austria) following the manufacturer’s protocol. It was sequenced with 56 other libraries in equimolar concentrations in one lane of an Illumina NovaSeq S4. Low quality reads were identified and removed using Trimmomatic (Bolger et al, 2014).

### Sequence assembly and annotation

The transcriptome assembly software Trinity v2.10.0 (Grabherr et al, 2011) was used to assemble the RNA-seq data into transcripts, with the Trimmomatic (Bolger et al, 2014) argument specified to trim sequences prior to assembly. TransDecoder v5.5.0 (github.com/transdecoder) was then used to predict putative coding regions within the transcripts for each assembly. Candidate coding sequences were first predicted using TransDecoder.LongOrfs. These sequences were screened against the Uniprot/SwissProt curated protein dataset using BLAST, along with the PFAM-A protein domain database using HAMMER, to obtain homology information to improve overall coding sequence prediction. Finally, TransDecoder.Predict was used to generate the final high-quality coding sequence predictions. The coding sequences for each assembly were then searched for 156 genes of interest using BLAST. A multi-fasta file containing the genes of interest was manually compiled by searching the *Alligator mississippiensis*, *Alligator sinensis*, *Protobothrops mucrosquamatus*, *Gallus gallus*, *Chrysemys picta*, *Anolis carolinensis*, *Python bivittatus*, *Pantherophis guttatus*, *Gekko japonicus*, *Columba livia*, *Serinus canaria*, *Aquila chrysaetos*, *Pseudopodoces humilis*, and *Cuculus canorus* (SI Table 6) gene annotations for the genes of interest. BLAST hits were filtered for the best hit using bitscore, while also ensuring that the subject and target sequence lengths were comparable. For each transcriptomic reference, the best hit genes were extracted into a multi-fasta file, which formed the reference files for the gene capture data.

Gene capture data was processed through a custom consensus pipeline based on the methods from Schott et al (2017). FastQC v0.11.9 (FastQC, 2015) was first used to assess the quality of each gene-capture sample. The data was then quality trimmed using Trimmomatic to remove adapter sequences and any low quality bases from the ends of reads. Gene capture samples were then aligned to one of the eleven reference files based on their evolutionary distance. The software BWA (Li and Durbin, 2009) was used to align the gene-capture samples to their respective reference files with parameters -B 2 to reduce the mismatch penalty (default value 4) and -M to mark split alignments as secondary. Unmapped reads were then removed from the alignment files using SAMtools (Li et al, 2009; Danacek et al, 2021). Genotyping was performed using the SAMtools/BCFtools mpileup and call pipeline (Li, 2011). SAMtools mpileup was run with parameters -d 5000 to increase the maximum depth, -Q 20 to control the mapping quality and -q 20 to control the base quality (Schott et al, 2017). Genotype calls were then converted to variant calls using BCFtools call. Variants were normalised using BCFtools norm, with multiallelic SNPs being joined into single records using -m +any. Finally, consensus sequences for each gene-capture sample relative to the reference genes of interest were generated using BCFtools consensus with argument -H 1 to use the first allele from the GT field.

We complemented this dataset with complete CDS regions of opsins for other squamates available on GenBank (SI Table 1). Additionally, full annotated genomes sourced from Genbank were BLASTed (E-value ≤ 0.05) for individual exons of opsin genes. BLAST hits were then curated and concatenated to construct full coding sequences. These were then aligned and any intron sections or UTRs still present were identified and deleted.

The opsin genes were aligned using Clustal Omega (Sievers et al, 2011) in Geneious Prime (v2020.2.4). We identified spectral sites based on Yokoyama, 2008a (and references therein), as well as other influential sites as listed in the entries on NCBI and Uniprot to determine the sites known to alter spectral sensitivity in the opsins. Leading into the analyses, 39 *Lerista* and 20 *Ctenotus* species from gene capture remained in the alignments. Opsin identity was confirmed by assembling a larger alignment combining (using MAFFT; Katoh et al, 2002) all the opsin sequences and generating a tree in order to verify the identifications of the opsins. Firstly, a nucleotide evolution model was selected using ModelTest-NG (Darriba et al, 2020) in raxmlGUI 2.0 (Edler et al, 2020). The chosen evolutionary model (JTT+G4+F) was then used to construct a tree with RAxML-NG (Koslov et al, 2019), also using raxmlGUI 2.0 (Edler et al, 2020). The outgroup was set to be the LWS opsin cluster (Terakita, 2005). This tree was then checked visually for opsin clustering to support their likely identity.

### Analyses of molecular evolution

Firstly, we downloaded reference opsin sequences from GenBank (SI Table 1), and aligned them with the skink gene captures in order to quickly identify any changes to important gene regions in skinks.

To detect changes in selection on skink opsin genes, CodeML analyses were conducted using PAML v4.8a (Yang, 2007) via ete3 (Huerta-Cepas et al, 2016). CodeML allows comparison of codon-based models of selection based on the ratio of non-synonymous (dN) to synonymous (dS) substitution rates (dN/dS, or ω). dN and dS are expected to occur at the same rate (ω ≈ 1) under a model of neutral evolution, whereas sites evolving under purifying (negative) selection are expected to have ω < 1, and those under diversifying (positive) selection are expected to have ω > 1 (Álvarez-Carretaro et al, 2023). However in tests such as the branch test, ω > 1 is not often seen as diversifying selection would not be expected across most sites (Álvarez-Carretaro et al, 2023). It is worth noting here that these CodeML models struggle to distinguish an increase in positive selection intensity versus relaxation of either purifying or positive selection (Wertheim et al, 2015). Therefore CodeML tests were supplemented by RELAX, which implements an alternative model with a selection intensity parameter *k.* If the addition of *k* improves model fit, the null model is rejected, with *k > 1* indicating intensified selection and *k < 1* indicating relaxation in the test partition (Wertheim et al, 2015).

To implement CodeML (Yang, 2007) a well-resolved skink species tree based on a multi-locus maximum likelihood analysis (Pyron et al, 2013) was used as the input tree in all analyses. This tree was reduced by pruning tips, retaining only the 95 skink species sampled here for opsin genes. Site models were performed on all skinks species, whereas branch, branch-site and clade models were performed only on *Lerista*. This is because *Ctenotus* do not differ in digit number or limb configuration, and didn’t differ in spectral tuning or opsin configuration along ecological lines.

We used site models - M0, M1a, M2a, M7 β, M8 β&ω to estimate the ratio of synonymous and non-synonymous substitution rates (ω) across amino-acid sites of the skink opsin genes. Likelihood ratio tests (LRTs) were used to compare the fit of null model M1a to the alternative model M2a, and null model M7 β to the alternative M8 β&ω. Bayes Empirical Bayes (BEB) was used to infer the sites under positive selection under the M2a and M8 β&ω models.

Branch models (one ratio model, two ratio model, free ratio model) allow ω to vary according to branches on the phylogenetic tree (see Fig. 1). These analyses were performed only for *Lerista* species because these show the highest ecological and morphological diversity. The groupings of *Lerista* were: climate, in which the 19 species found in arid and semi-arid habitats (hereafter termed ‘*arid*’ group) were the foreground and 20 species that occupy tropical and temperate habitats (hereafter ‘*temperate-tropical*’ group) were the background; eyelid condition, where 7 species with fused eyelids (‘*fused*’ group) were the foreground and 32 with unfused eyelids (‘*unfused*’) the background; and limb condition, where 27 species with <3 hindlimb digits were in the foreground (*<3 hindlimb digits*) and 12 other species were in the background (*≥3 hindlimb digits*, see Fig. 1). Limb configuration information was obtained from Cogger (2014).

Branch-site models (model A, model A1, model B) allow ω to vary across both sites and branches and this was used to determine if positive selection was present at certain sites and differed according to the ecological or morphological groupings of *Lerista*. The null model to model A is model A1, where ω2=1 as according to Zhang et al (2005).

Clade analyses (CmC, CmD) have site classes such that ω<1, ω=1 and 0<ω<∞, and assume that some sites evolve conservatively across the phylogeny whereas a class of sites can evolve freely. Analyses were performed between *Lerista* and *Ctenotus* groups. The null model for CmC is M2a_rel (Weadick and Chang, 2012), which constrains CmC such that ω2=ω3.

Due to difficulties in distinguishing positive and relaxed selection (Wertheim et al, 2015), we also conducted tests using the RELAX analysis on Datamonkey.com (Wertheim et al, 2015; Weaver et al, 2018). This tests the gene-wide strength (intensification versus relaxation) of selection on user selected branches of a given tree using the given alignments. Firstly, test and reference branches are selected by the user, then a null model is compared to an alternative model, where the null model has a selection intensity parameter (k) = 1, and the alternative has k as a free parameter. An LRT then determines if the alternative model has better fit to the data than the null model. If so, this indicates a significant difference in selection intensity on the test branch compared to the reference branch, with k<1 indicating relaxation of selection and k>1 indicating intensification of selection. We used the same trees, alignments and branch selections as in the branch and branch-site tests (see above). We also performed a control on branch, branch-site and clade analyses. We performed M1a and M2a analyses on background branches only. This was done because codeml assumes neutral or purifying selection on background branches. If positive selection was found, this invalidates any significant differences that branch, branch-site and clade analyses found between the foreground and background branches.

In all M2a and M8 β&ω site analyses, and branch site analyses, sites potentially under positive selection were detected by the Bayes Empirical Bayes method with posterior probabilities of >0.95 (Yang et al, 2005). Any selected sites were reported using the amino-acid number associated with *Bos taurus* RH1.

## Results

### Eye size

Fig. 3 shows the variation in relative eye size in *Lerista* and *Ctenotus*. In both genera eye size diminished in log linear fashion with reduced head size, but whereas relative eye size remained tightly constrained in *Ctenotus*, in *Lerista* relative size was more variable and the data fell into roughly two groups on either side of the overall trend line. Above the line were species and clades in which fore and hind limbs were retained, mostly with three or more digits on each limb. Below the line were species and clades with greater limb reduction and body elongation, with the forelimbs either missing or greatly reduced. *Lerista apoda*, the most extreme in terms of eye size reduction, also has the greatest degree of body elongation (mean presacral number over 60, Greer 1987), is one of only two species with no external trace of any limbs, and is the only species in which the eye is buried under a cephalic scale rather than enclosed by (moveable or fixed) eyelids.

**Figure 3:**
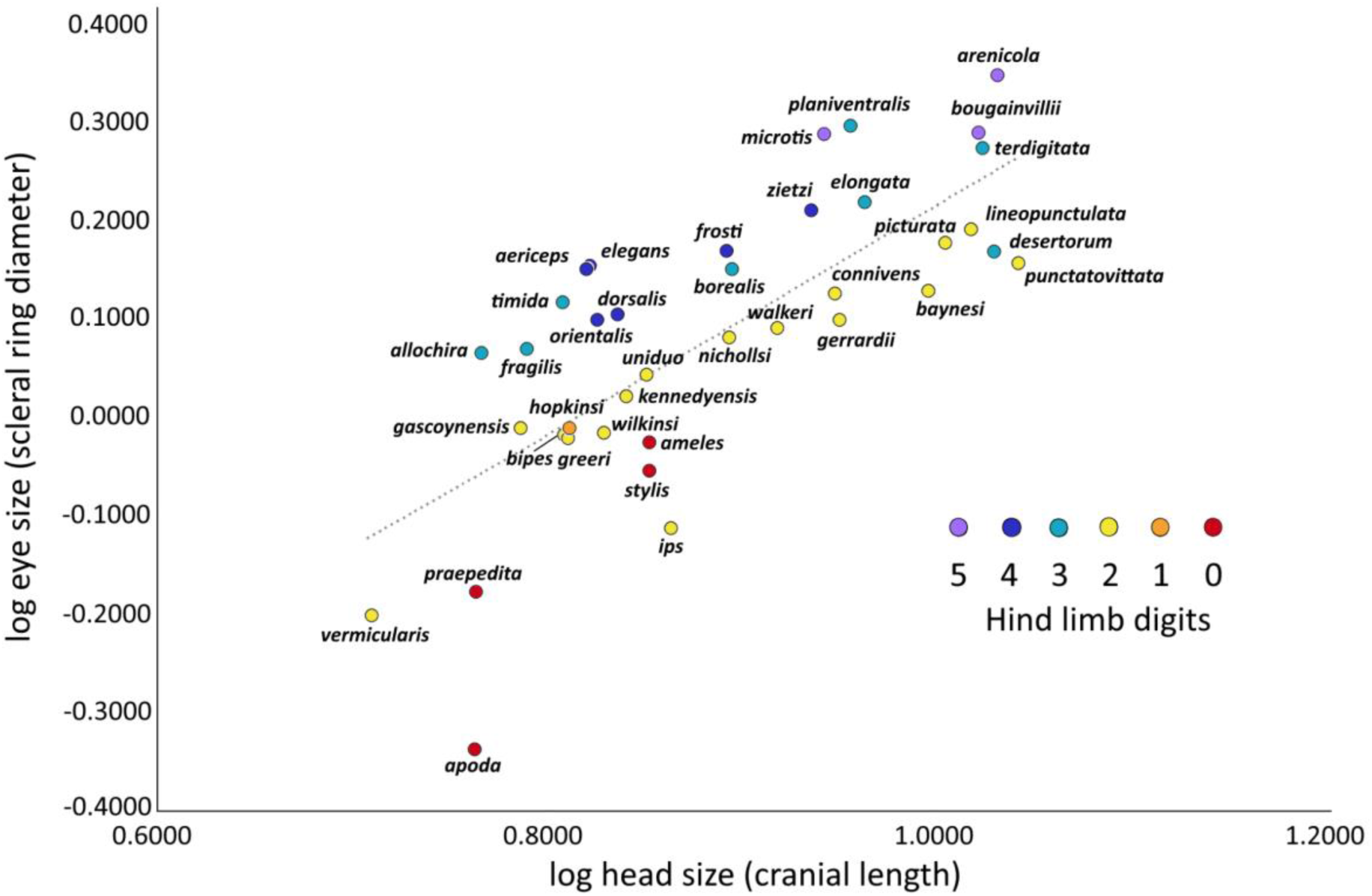
Comparison of eye size to head length in sampled Lerista species, also showing the number of hind limb digits in each species and showing a clear relationship between eye/head size ratio and hind digit number. The majority of species with ≥3 hindlimb digits have an eye size greater than ∼17% of head length, while most species with <3 hindlimb digits have an eye size less than ∼17% of head length.

### Targeted capture of skink opsins

From a total of 92 samples, full length sequences were obtained from 87 species for LWS, 88 for SWS1, and partial sequences were obtained from 85 species for RH1 (630bp), 95 for RH2 (492bp) and 87 for SWS2 (438bp) (SI Table 1). All opsin genes were found in 59 *Lerista* and *Ctenotus* sequenced. Each opsin resolved as a monophyletic grouping (SI Fig. 6). No gene duplications were found. We found some full sequences in closely related species, so we feel confident in reporting these sequences are retained in our study species, just not fully captured. *Lerista zietzi* and *L. timida* yielded low quality sequences that were discarded. From transcriptomes, full sequences of each 12 of 13 sequences were found in LWS, with *Melanoseps occidentalis* being near complete. In RH1, 11 partial sequences were found, again with *Melanoseps occidentalis* being near complete. RH1 was not found in *Eutropis macularia* and *Tiliqua rugosa*. In RH2, 11 sequences were found, with one full sequence in *Platysaurus broadleyi*, and near complete sequences in *Chalcides ocellatus*, *Eumeces schneideri*, *Eutropis macularia*, and *Glaphyromorphus punctulatus,* with six partial sequences for other species. RH2 was not found in *Melanoseps occidentalis* and *Tiliqua rugosa.* In SWS1, 12 sequences were recovered. 10 were complete, while *Eutropis macularia* and *Melanoseps occidentalis* were near complete. SWS1 was not found in *Tiliqua rugosa*. In SWS2, 12 sequences were recovered. Full sequences were found in *Chalcides ocellatus* and *Platysaurus broadleyi*, and partial sequences were found in the other species. SWS2 was not found in *Tiliqua rugosa* (SI Table 1).

One limiting factor in this study is the fact that RH1, RH2 and SWS2 were not recovered as full sequences for any of the gene captures conducted. SWS2 and RH2 recovered here in particular are quite short (438bp and 492bp respectively), at less than half of their complete size (1092bp and 1068bp respectively in *Anolis carolinensis*). This means many known tuning sites on these opsins are not represented in this data (7 for RH1, 2 for RH2, 4 for SWS2 were recovered), and thus the potential variance within cannot be examined. Therefore we must estimate that the sensitivity of the opsins of *Lerista* and *Ctenotus* are similar to previously described skink species (*Tiliqua rugosa*) and are as follows: LWS ≈ 560nm; RH1 ≈ 491nm; RH2 ≈ 495nm; SWS1 ≈ 360nm and SWS2 ≈ 440nm (Nagloo et al, 2016). Further evidence that these regions are conserved in *Lerista* and *Ctenotus* comes from the recently released genome of *Lerista edwardsae* (accession number GCA_029204185.1), where the full SWS2 and RH2 genes can be recovered. Were the full sequences present in this study, we would have a much clearer picture of spectral tuning in all opsins and could potentially discover variances at known spectral tuning sites.

### Analyses of molecular evolution

Site models show strong signals of positive selection at the codon level in four of the five opsins; only RH1 exhibits weak evidence of positive selection (Table 1; SI Table 2).

**Table 1:** Significantly selected sites according to BEB under site-models. Amino-acid numbering based on the respective position in the bovine rhodopsin.

| Models | Sites Under Positive Selection (BEB) |
| --- | --- |
| ***sws1* opsin gene** |  |
| M8 ($\beta&\omega$) | 50 |
| ***lws* opsin gene** |  |
| M2a | 39 - 112 - 155 - 217 - 270 |
| M8 ($\beta&\omega$) | 39 - 112 - 155 - 162 - 217 - 262 - 270 |
| ***rh2* rhodopsin gene** |  |
| M8 ($\beta&\omega$) | 304 |
| ***sws2* rhodopsin gene** |  |
| M2a | 11 - 88 - 108 - 116 |
| M8 ($\beta&\omega$) | 11- 88 - 89 - 108 - 116 - 123 |

However, most branch model tests did not find statistically significant differences in selection pressures between partitions. These results are described for each gene below.

#### Site models

The LWS gene shows evidence of positive selection at seven amino acid sites. Models M2a and M8 β&ω are a significantly better fit than their respective null models, M1a and M7β. According to BEB for M2a and M8 β&ω, sites five sites in helical transmembrane domains, and one in the first extracellular loop (Fig. 4) are under positive selection (sites 162 and 262 (both in transmembrane regions) are inferred to be under positive selection under M8 β&ω (but not under M2a). None of these sites are known to have an impact on spectral tuning of LWS, but sites 262 and 270 are one amino acid directly after a spectral tuning site (277 and 285 by bovine LWS), site 162 is 2 amino acids from a spectral tuning site (180 by bovine LWS) and site 112 is the amino acid directly before a retinal chromophore binding site (this is in the extracellular loop in bovine LWS but in a transmembrane region in bovine RH1)(Table 1; SI Table 2).

**Figure 4:**
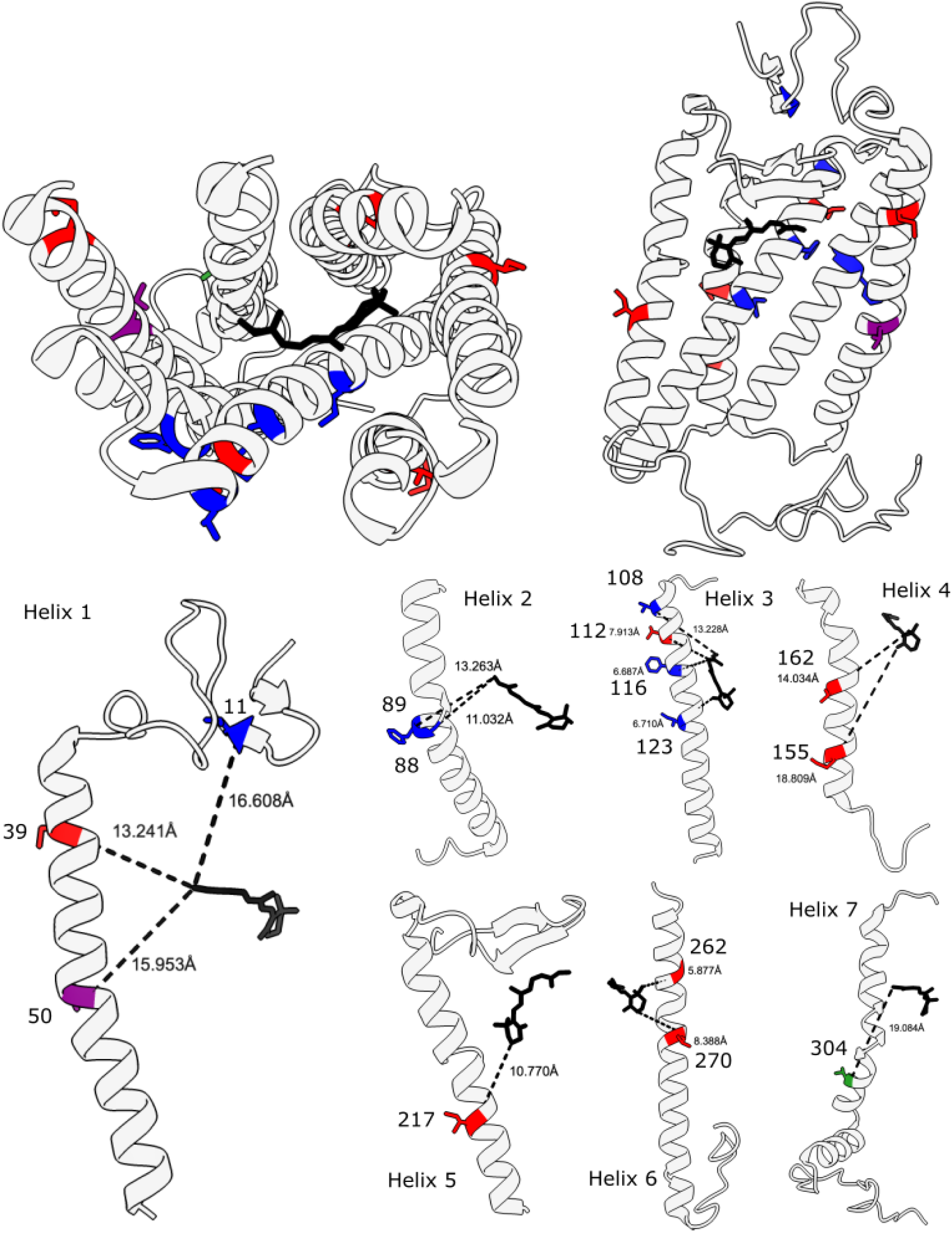
Depiction of bovine RHO (1u19) with positively selected sites found in this study shown as follows: LWS, red; SWS1, purple; SWS2, blue; RH1, green. The chromophore is shown in black. The left image has had the extracellular portions removed for better clarity. The right image has had helices 6 and 7 removed for better clarity

The RH1 gene shows evidence of positive selection in two amino acid sites (SI Table 2). Models M2a and M8 β&ω are not a significantly better fit than their respective null models, M1a and M7 β, however M7 β/M8 β&ω is not far from significance (p=0.081). According to Bayes Empirical Bayes (BEB) for M8 β&ω, sites 201 and 219 are under positive selection (Fig. 4). Neither of these sites are known to have an impact on spectral tuning of RH1, but site 201 is at a zinc metal binding site.

The SWS1 gene shows evidence of positive selection at one amino acid site. Model M8 β&ω is a significantly better fit than its null model M7 β, but M2a was not a significantly better fit than M1a. According to Bayes Empirical Bayes (BEB) for M8 β&ω, site 50 (transmembrane) is under positive selection (Fig. 4). M2a did not show any selected sites. Site 50 is not known to have an impact on spectral tuning of SWS1, but it is the amino acid directly after a spectral tuning site (site 49) (Table 1; SI Table 2).

The SWS2 gene shows evidence of positive selection in six amino acid sites. Models M2a and M8a are a significantly better fit than their respective null models, M1a and M7 β. According to Bayes Empirical Bayes (BEB) for M2a and M8 β&ω, two sites in transmembrane regions and two sites in extracellular regions are under positive selection (Fig. 4) (sites 89 (transmembrane) and 123 (transmembrane, retinal chromophore binding pocket) are inferred to be under positive selection under M8 β&ω but not M2a) (Table 1; SI Table 2). None of these sites are known to have an impact on spectral tuning of SWS2.

The RH2 gene shows positive selection at one amino acid site. Models M2a and M8 β&ω are a significantly better fit than their respective null models, M1a and M7 β. According to Bayes Empirical Bayes (BEB), only site 304 (transmembrane) is under positive selection (Fig. 4); however this is only under M8 β&ω but not M2a. This site is not known to have an impact on spectral tuning of RH2 (Table 1; SI Table 2).

#### Branch Models

In LWS, branch models the limb condition analysis produced a significant difference between groups (p=0.0393), with the <3 hindlimb digits group having a higher ω (ω=0.151) than the ≥3 hindlimb digits group (ω=0.0664). All other groups did not produce significant differences. Giving each branch its own ω did not produce significantly different results to all branches having a single fixed ω (SI Table 3). For group comparisons in all other visual opsin genes, no test produced a significant difference. Giving each branch its own ω did not produce significantly different results to all branches having a single fixed ω (SI Table 3).

#### Branch-Site Models

In LWS, branch-site models reveal that for the arid climate group and *<3 hindlimb digits*, site 112 (Fig. 4) is under positive selection. For *<3 hindlimb digits,* site 270 is also under selection. No other branch-site analyses revealed significant results (SI Table 4). For RH1 and SWS1, branch-site models reveal that no sites were significantly selected for in any group.

For SWS2, branch-site models reveal that for the *arid* group, two sites were under positive selection. For *<3 hindlimb digits*, six sites were under positive selection. There was no evidence for positive selection at any site in the eyelid analysis (SI Table 4).

For RH2, branch-site models reveal that for the arid climate group, four sites were under positive selection. For *<3 hindlimb digits*, three sites are under positive selection. There was no evidence for positive selection at any site in the eyelid analysis (SI Table 4)

#### Clade Models

For LWS, in clade analyses, CmC is a significantly better fit to data than M2a_rel (SI Table 5). In the eyelid analysis, *mobile* (ω=0.00723) versus *fused* (ω=0) groups. The ω values were lower in *temperate-*tropical (ω=0) versus arid climate groups (ω=0.009). Similarly, <3 hindlimb digits group had a higher ω (ω=0.009) than the ≥3 hindlimb digits group (ω=0) (SI Table 5).

For RH1 in clade analyses, CmC is a significantly better fit to data than M2a_rel (SI Table 5). The *fused* vs *mobile* eyelids analysis showed significant site-specific positive selection in both partitions (SI Table 5).

For SWS1, SWS2 and RH2 clade analyses, CmC is a significantly better fit to data than M2a_rel for all groups, but no evidence of site specific positive selection was found (SI Table 5).

#### RELAX Tests

LWS was also the only gene that passed the significance cut-off for model testing using RELAX. In this gene, the test branches with *<3 hindlimb digits* showed a significant relaxation of the intensity of selection (K=0.73; p=0.014) relative to the reference branches (*≥3 hindlimb digits* group) (SI Table 7).

Drop-out analyses indicated that all background groups of RH2 and SWS2, climate and eyelids in LWS, and eyelids in SWS1 and RH1, all had evidence of positive selection. Therefore the conclusions of differences between the foreground and background branch, branch-site and clade tests as described previously must be rejected.

## Discussion

Our study provides a detailed view of the molecular evolution of visual opsins across a recent radiation of 126 ecologically diverse scincid -+lizards (including 39 *Lerista* and 20 *Ctenotus* species, 67 other Scincoidea). Sequence analyses of full-length SWS1 and LWS and (mostly) partial SWS2, RH1 and RH2 sequences suggested that these visual opsins have been retained - no stop codons or frameshift mutations were observed in any of the sampled species. Consistent with this, we detected significant signals of positive selection in four of the five opsins (LWS, SWS1, SWS2, RH2). Most comparisons of site-, branch- and clade-specific selection pressures did not show significant differences according to broad ecological divisions between lineages inhabiting arid versus temperate-tropical climates, and lineages with diurnal surface-active habits versus highly fossorial forms with small, recessed eyes or fused eyelids. This along with the data about eye size relative to body size provides evidence that evolution of fused eyelids and spectacles is not correlated to limb or digit loss and burrowing behaviour, rather that it evolves most reliably in species with small body size but larger relative eye size. These data support Greer’s (1983) hypothesis that the spectacle tends to evolve in small-sized species, and may be a response to preventing water loss via the cornea. Most of those with fused eyelids ("spectacle") are among those with the largest relative eye size, but all are among the smaller-bodied species. Of those species with head length less than the sample mean (7.76 mm or less), 12 of the 15 larger-eyed species ("above the line") have a spectacle (exceptions are *Lerista viduata*, *L. fragilis* and *L. dorsalis*), while 7 of 17 smaller-eyed species have a spectacle (Fig. 3). The one area where a spectacle is starting to look like a correlate of eye reduction is just the two smallest-eyed species, *L. vermicularis* and *L. apoda*, which are the extremes in *Lerista* smallness (Greer, 1983). A significant signal of relaxed selection pressure was detected only in the LWS of limb reduced versus fully limbed *Lerista,* which is an unusual occurrence, as LWS is normally retained in ecological transitions (Jacobs, 2013; Sadier et al, 2018; David-Gray et al, 2002; Davies, 2011, Simões et al, 2019). The retention and ongoing positive selection on visual opsins, particularly cone opsins associated with bright-light vision, in these ecologically diverse species has important implications for understanding the impact of photic transitions on visual function and evolution in lizards.

### Conservation versus loss of visual genes in cryptozoic squamates

Our results provide an important contribution to our understanding of visual evolution during ecological transitions in squamates. Previous studies of visual characteristics in fossorial and dim-light adapted snakes (Simões et al, 2015, 2016; Gower et al, 2021), and geckos (Pinto et al, 2019) have revealed multiple opsin losses. Estimates of the age of these lineages, and therefore the maximum time these losses have had to occur, are around 120 mya for Scolecophidia (Miralles et al, 2018), and 112 mya for the extant gekkotans (Daza et al, 2012; 2014), with the origin of nocturnality being close to the root of Gekkota (Anderson et al, 2017; Gamble et al, 2015). However, other fossorial squamates, of similar evolutionary age, show retention of vision genes and visual capabilities. In particular, the amphisbaenian lizards, which diverged from other Lacertoidea ∼138mya (Vidal and Hedges, 2005), are highly fossorial and have reduced eyes yet have retained and express a full complement of visual opsins (Simões et al, 2015). This may mean that eye size might be reduced in taxa where re-modelling head shape for fossorial behaviour is occurring, but not necessarily that photopic function is being lost. This could also be true for other limb-reduced lizard clades. These variable patterns of opsin retention and loss in scotopic squamates prompt the question of whether the *Lerista* lineages analysed here are simply too recent to have accumulated inactivating mutations in their opsin sequences. Based on fossil-dated molecular relationships, *Lerista* underwent several independent transitions from surface active to fossorially adapted forms over the last 13.4 million years (Skinner et al, 2008). There is limited comparative data on the rate of visual gene loss during ecological shifts in most taxa. However, evolutionary regression is expected to proceed most quickly if this is driven by selection favouring e.g. energy saving, while longer timeframes are expected if degeneration is a consequence of genetic drift due to lack of function (Jeffery, 2009). In two species of cavefish, *Lucifuga dentata* and *Astyanax mexicanus,* relaxation in pseudogenised vision genes is estimated to have have taken from 1.3-1.5mya, or 370,000 generations, to as little as 20,000 years ago (Policarpo et al, 2021; Moran et al, 2015; Fumey et al, 2018). In neotropical cichlids, many species have lost blue sensitive opsin genes (Hauser et al, 2021), and where most clades that have undergone these losses are >90my old, *Uaru fernandezyepezi* may have diverged from its closest relative ∼15mya (López-Fernández et al, 2012). This is around the age of *Lerista* (Skinner et al, 2008), with the loss of all limbs from species such as *L. ameles*, *L. apoda* and *L. stylis* having occurred in a period of 11.8my (Skinner et al, 2008). *Lerista* species that move headfirst through solid substrates might also be expected to experience adaptive selection for eye size reduction; eyes are naturally delicate instruments and reducing their size obviously reduces the potential of damage. However, lizards that have periods of activity in well-lit microhabitats on or near the surface would be expected to retain eye function, which may be why *Lerista* does not appear to have experienced much direct selection for opsin regression (See SI Tables 3 and 7). The fastest rate of limb loss from pentadactyl or tetradactyl to adactyl may have occurred in as little as 3.6my (Skinner et al, 2008), indicating that losses of major anatomical structures in these animals have occurred very quickly; that major loss of retinal function has not occurred indicates a continuing utility in spite of reduction in eye size.

Relaxation of purifying selection in the LWS of limbless and limb reduced *Lerista* suggests that detection of longer wavelengths is not as tightly constrained in these lizards compared to fully limbed *Lerista*, indicating somewhat smaller reliance (though far from relaxed as it still under purifying selection) on long wavelength detection. This is very unexpected given that very few organisms have been reported to have lost this opsin previously. Ordinarily, fossorial species tend to show spectral shifts to longer wavelengths, and if any opsins are lost or pseudogenised, LWS is often the least likely candidate (Peichl et al, 2001; David-Gray et al, 2002; Jacobs, 2013; Simões et al, 2019). In many species across the animal kingdom, fossorial or nocturnal species, or species with scotopic visual traits, lose function in opsins tuned to shorter wavelengths, maintaining LWS (Jacobs, 2013; Sadier et al, 2018; David-Gray et al, 2002; Davies, 2011, Simões et al, 2019). For instance, the obligate fossorial mole rats possess subcutaneous eyes and are unable to respond to visual images, but still maintain LWS monochromacy for photoentrainment (David-Gray et al, 2002). Among vertebrates, the most notable losses of LWS have been in deep sea fish (Musilova et al, 2019) and deep diving whales (Meredith et al, 2013; Schweikert et al, 2016; Springer et al, 2016), which occupy photic habitats dominated by shorter wavelengths (Warrant and Johnsen, 2013), thus have little ecological parallel with our study system. A more detailed understanding of the visual ecology of *Lerista* may shed light on the cause and significance of increased relaxed selection on their LWS opsin gene.

### The link between visual function and eye-reduction in fossorial skinks

Absolute eye size is known to affect several aspects of visual function, not least of which is visual acuity (Walls, 1942). As eye diameter increases, and assuming all other eye parameters are constant, posterior nodal distance (PND) increases and creates a larger retinal image (Veilleux and Kirk, 2014; Hughes, 1977). Walls (1942) suggests that because of the higher sampling by photoreceptors, visual acuity should be increased, and various studies have since provided some support for this in mammals (Kirschfeld, 1976; Veilleux and Kirk, 2014). In smaller eyes, photoreceptors can narrow in order to make up for the loss of acuity due to shorter focal length, however this also means lower light collection per photoreceptor and thus there is a trade off between acuity and sensitivity (Caves et al, 2018).

It is important to note, we do not know the extent to which these lizards are diurnal and surface active due to their cryptozoic habits. We lack species specific information on visual ecology with which to make robust links between visual and molecular adaptation. It has been suggested that the great majority of *Lerista* species will also have periods of diurnal activity as well as crepuscular and nocturnal (Meiri, 2018). Any difference in their overall exposure to light and light conditions will be a difference of degree rather than of kind. It should also be noted that although *Lerista* do possess many adaptations associated with a heavily fossorial lifestyle, they also all possess an external ear (Greer, 1967), something often lost in heavily fossorial squamates. They also all possess a dark peritoneum, the abdominal cavity lining (Hutchinson, personal observation), which has been associated with diurnal activity (Greer, 2002).

### Adaptive opsin evolution during the *Ctenotus-Lerista* radiation

The results of the site model analysis seem to show that there is indeed selection at many sites, and some of the variation at these sites can be explained by ecological categories according to branch site models. Climate and limb analyses produced significant results in LWS, RH2 and SWS2 (SI Table 4). Without biochemical analyses, we cannot know the significance of these changes, whether they impact spectral sensitivity, or to what extent. Many of the sites listed as positively selected in this study seem to be close to previously known tuning sites. It is well known that changes at tuning sites can significantly alter the peak spectral sensitivity of the photopigment, but it is less clear how any changes in other positions along the opsin might affect tuning, if at all. There are a few changes that seem to happen mostly in conjunction with one another and occur in close proximity to one another (sites 97 and 98, SWS2 most notably). The phenomenon of previously undescribed positively selected sites being found in close proximity to previously known ones has also recently been described in anurans (Schott et al, 2022). Perhaps here too, these previously unidentified and unexplored sites may have minor effects on spectral tuning or other functions of the opsin, such as retinal release and binding, rates of retinal bleaching and recovery and so on.

One of the most interesting findings of the site analyses is the selected site at 201 on RH1. Although the site did not quite meet probability thresholds (p<0.05), there is variation at this site, and skinks as a group seem to have diverged from other groups. This site has been previously identified as a zinc binding site (Toledo et al, 2009; Norris et al, 2021). Zinc is known to have a number of binding sites on rhodopsin, with site 201/279 possibly being a non-specific one (Toledo et al, 2009). The zinc binding sites increase the stability of the pigment, especially between the first extracellular loop and the sixth helix (Park et al, 2007). Its sister site, site 279, is consistent across all skinks, *Anolis* and *Python* and has the same amino acid as *Bos*, Glutamine. However site 201 seems to differ in skinks, and is not consistent throughout the clade, with only *Liopholis striata* possessing Glutamic acid at this site. Other skinks possess either Arginine and Lysine, with only *Plestiodon fasciatus* possessing a different amino acid, Glutamine. None of these amino acids are one of the four that have been found to commonly bind to zinc (Shu et al, 2008). Therefore it would appear that this zinc binding site may have been made non-functional across Scincidae. Interestingly, *Sphenodon* genome shows Lysine at this site too. The observation that this site seems to have changed frequently in skinks (and *Sphenodon*), yet is conserved in other clades in Squamata and even Mammalia, suggests a strong evolutionary advantage in decreasing the stability of this opsin. The other sites which were found to be under selection in this study also fall into the region that is particularly destabilised by the absence of zinc binding (Park et al, 2007).

Although the exact effects of the loss of this one particular zinc binding site are unclear, one may speculate that it would have a destabilising effect, and perhaps accentuate any spectral tuning and tertiary structure changes that the selected sites could cause. Helices 5 and 6 are known to undergo conformational changes when rhodopsin is activated (Zhou et al, 2012) and destabilisation of this region by the removal of zinc binding may have implications for the activation and function of rhodopsin. Further study is needed to understand the functional significance of these findings.

## Conclusions

This study used thorough taxon sampling spanning replicate ecological transitions to investigate visual changes in skinks. Losses of cone opsins may have been expected in the most specialised fossorial lineages, as happened in geckos and snakes in historical low light transitions. Instead, all five opsins are retained in even the most dedicated fossorial lineages. We suggest that visual systems in these species have retained adaptive value in habitats with reduced visible distance. The *Lerista* clade maintains its status as a key system for low light adaptive studies, but the results of this study suggest that the variations and extremity of adaptations are much more subtle than previously described, and more research is needed to understand the species’ activity patterns and extent of their fossorial activity. Further work on the particular effects of changes to the zinc binding sites, and the potential implications this may have for spectral tuning in the destabilised regions, would also help to shed light on the significance of the findings of this paper. Finally, we recommend morphological studies to examine a possible link between the patterns of gene selection reported here and fossorial adaptations of traits including skull shape and eye size.

## Author Contributions

M.J.R.F., K.L.S. and B.F.S. conceived the study. B.F.S. and M.N.H. conducted fieldwork. M.J.R.F. and B.F.S. did the laboratory work with the support of T.B. Bioinformatic analysis were done by M.J.R.F. with the support of A.J.L. Phylogenetic and comparative genomic analyses were performed by M.J.R.F. with input from B.F.S., M.N.H. and K.L.S. Results were interpreted by M.J.R.F., B.F.S. and K.L.S. with input of M.N.H. Morphological analysis was done by M.N.H. The manuscript was written by M.J.R.F. with support of B.F.S. and K.L.S. and with input from all coauthors.

## Supplementary Materials

Table 1. List of specimens with SA Museum ABTC numbers, NCBI numbers, type of sequence and presence of each visual opsin

Table 2. Site CodeML results

Table 3. Branch CodeML results

Table 4. Branch site CodeML results

Table 5. Clade CodeML results

Table 6. Genes Sampled with Probes and Probe Used

Table 7. RELAX Results

## Acknowledgements

In addition to thanking all co-authors for their tireless work on this study, I would like to thank the South Australian Museum for allowing access to and use of their tissue collections. Suggestions from Einat Hauzman and Fabio Cortesi also greatly improved this manuscript.

