## Supplemental Tables for "Retention of the full visual opsin repertoire in Australia’s cryptozoic lizards"

Table 1. List of specimens with SA Museum ABTC numbers, NCBI numbers, type of sequence and presence of each visual opsin

| Species | ABTC Number | NCBI number | Type of Sequence | LWS | RH1 | RH2 | SWS1 | SWS2 |
|---|---|---|---|---|---|---|---|---|
| Chalcides ocellatus | NHM_E36 | | Gene captures | ✓ | ✓ | ✓ | ✓ | ✓ |
| Corucia zebrata | 50360 | | Gene captures | ✓ | ✓ | ✓ | ✓ | ✓ |
| Ctenotus pantherinus | 35092 | | Gene captures | ✓ | ✓ | ✓ | ✓ | ✓ |
| Ctenotus spaldingi | 57114 | | Gene captures | ✓ | ✓ | ✓ | ✓ | ✓ |
| Cyclodomorphus gerrardii | 11496 | | Gene captures | ✓ | ✓ | ✓ | ✓ | ✓ |
| Egernia striolata | 96398 | | Gene captures | ✓ | ✓ | ✓ | ✓ | ✓ |
| Emoia acrocostata | 50439 | | Gene captures | ✓ | ✓ | ✓ | ✓ | ✓ |
| Eugongylus rufescens | 98675 | | Gene captures | ✓ | ✓ | ✓ | ✓ | ✓ |
| Eulamprus quoyii | 85427 | | Gene captures | ✓ | ✓ | ✓ | ✓ | ✓ |
| Lampropholis guichenoti | 23334 | | Gene captures | ✓ | ✓ | ✓ | ✓ | ✓ |
| Lerista arenicola | 40786 | | Gene captures | ✓ | ✓ | ✓ | ✓ | ✓ |
| Lerista bougainvillii | 94931 | | Gene captures | ✓ | ✓ | ✓ | ✓ | ✓ |
| Lerista desertorum | 94330 | | Gene captures | ✓ | ✓ | ✓ | ✓ | ✓ |
| Lerista dorsalis | 106178 | | Gene captures | ✓ | ✓ | ✓ | ✓ | ✓ |
| Lerista edwardsae | 139775 | | Gene captures | ✓ | ✓ | ✓ | ✓ | ✓ |
| Lerista ips | 91595 | | Gene captures | ✓ | ✓ | ✓ | ✓ | ✓ |
| Lerista timida | 39962 | | Gene captures | ✓ | ✓ | ✓ | ✓ | ✓ |
| Liopholis inornata | 100983 | | Gene captures | ✓ | ✓ | ✓ | ✓ | ✓ |
| Liopholis whitii | 68829 | | Gene captures | ✓ | ✓ | ✓ | ✓ | ✓ |
| Liopholis striata | 91720 | | Gene captures | ✓ | ✓ | ✓ | ✓ | ✓ |
| Ophioscincus ophioscincus | 32202 | | Gene captures | ✓ | ✓ | ✓ | ✓ | ✓ |
| Plestiodon fasciatus | 12334 | | Gene captures | | ✓ | ✓ | ✓ | ✓ |
| Pseudemoia entrecasteauxii | 23134 | | Gene captures | ✓ | ✓ | ✓ | ✓ | ✓ |
| Saiphos equalis | 14195 | | Gene captures | ✓ | ✓ | ✓ | ✓ | ✓ |
| Saproscincus challengeri | 11015 | | Gene captures | ✓ | ✓ | ✓ | ✓ | ✓ |
| Silvascincus silvascincus | 12360 | | Gene captures | ✓ | ✓ | ✓ | ✓ | ✓ |
| Sphenodon puncatatus | 32244 | | Gene captures | ✓ | ✓ | ✓ | ✓ | ✓ |
| Tiliqua scincoides | 57724 | | Gene captures | ✓ | ✓ | ✓ | ✓ | ✓ |
| Anomalopus brevicollis | 10849 | | Gene captures | ✓ | ✓ | ✓ | ✓ | ✓ |
| Anomalopus gowi | 10861 | | Gene captures | ✓ | ✓ | ✓ | ✓ | ✓ |
| Anomalopus leuckartii | 53604 | | Gene captures | ✓ | ✓ | ✓ | ✓ | ✓ |
| Anomalopus pluto | 10961 | | Gene captures | ✓ | ✓ | ✓ | ✓ | ✓ |
| Anomalopus swansoni | 6970 | | Gene captures | ✓ | ✓ | ✓ | ✓ | ✓ |
| Anomalopus verreauxii | 138983 | | Gene captures | ✓ | ✓ | ✓ | ✓ | ✓ |
| Ctenotus agrestis | 113815 | | Gene captures | ✓ | ✓ | ✓ | ✓ | ✓ |

| | | | | | | | |
|---|---|---|---|---|---|---|---|
| Ctenotus allotropis | 8980 | Gene captures | ✔ | ✔ | ✔ | ✔ | ✔ |
| Ctenotus arcanus | 137937 | Gene captures | ✔ | ✔ | ✔ | ✔ | ✔ |
| Ctenotus ariadnae | 706 | Gene captures | ✔ | ✔ | ✔ | ✔ | ✔ |
| Ctenotus atlas | 10488 | Gene captures | ✔ | ✔ | ✔ | ✔ | ✔ |
| Ctenotus brevipes | 72811 | Gene captures | ✔ | ✔ | ✔ | ✔ | ✔ |
| Ctenotus brooksi | 133010 | Gene captures | ✔ | ✔ | ✔ | ✔ | ✔ |
| Ctenotus burbidgei | 105767 | Gene captures | ✔ | ✔ | ✔ | ✔ | ✔ |
| Ctenotus capricorni | 105818 | Gene captures | ✔ | ✔ | ✔ | ✔ | ✔ |
| Ctenotus coggeri | 30198 | Gene captures | ✔ | ✔ | ✔ | ✔ | ✔ |
| Ctenotus decaneurus | 29731 | Gene captures | ✔ | ✔ | ✔ | ✔ | ✔ |
| Ctenotus dux | 91591 | Gene captures | ✔ | ✔ | ✔ | ✔ | ✔ |
| Ctenotus essingtonii | 29141 | Gene captures | ✔ | ✔ | ✔ | ✔ | ✔ |
| Ctenotus euclae | 118396 | Gene captures | ✔ | ✔ | ✔ | ✔ | ✔ |
| Ctenotus eutaenius | 77199 | Gene captures | ✔ | ✔ | ✔ | ✔ | ✔ |
| Ctenotus gagudju | 29095 | Gene captures | ✔ | ✔ | ✔ | ✔ | ✔ |
| Ctenotus gemmula | 62119 | Gene captures | ✔ | ✔ | ✔ | ✔ | ✔ |
| Ctenotus grandis | 14303 | Gene captures | ✔ | ✔ | ✔ | ✔ | ✔ |
| Ctenotus greeri | 91836 | Gene captures | ✔ | ✔ | ✔ | ✔ | ✔ |
| Ctenotus hebetior | 9065 | Gene captures | ✔ | ✔ | ✔ | ✔ | ✔ |
| Ctenotus impar | 53530 | Gene captures | ✔ | ✔ | ✔ | ✔ | ✔ |
| Ctenotus ingrami | 32120 | Gene captures | ✔ | ✔ | ✔ | ✔ | ✔ |
| Ctenotus joanae | 56695 | Gene captures | ✔ | ✔ | ✔ | ✔ | ✔ |
| Ctenotus labillardieri | 58013 | Gene captures | ✔ | ✔ | ✔ | ✔ | ✔ |
| Ctenotus lancelini | 63038 | Gene captures | ✔ | ✔ | ✔ | ✔ | ✔ |
| Ctenotus leonhardii | 10011 | Gene captures | ✔ | ✔ | ✔ | ✔ | ✔ |
| Ctenotus militaris | 29855 | Gene captures | ✔ | ✔ | ✔ | ✔ | ✔ |
| Ctenotus mimetes | 82597 | Gene captures | ✔ | ✔ | ✔ | ✔ | ✔ |
| Ctenotus monticola | 105816 | Gene captures | ✔ | ✔ | ✔ | ✔ | ✔ |
| Ctenotus olympicus | 35032 | Gene captures | ✔ | ✔ | ✔ | ✔ | ✔ |
| Ctenotus orientalis | 119442 | Gene captures | ✔ | ✔ | ✔ | ✔ | ✔ |
| Ctenotus pantherinus | 137975 | Gene captures | ✔ | ✔ | ✔ | ✔ | ✔ |
| Ctenotus piankai | 23830 | Gene captures | ✔ | ✔ | ✔ | ✔ | ✔ |
| Ctenotus quinkan | 105817 | Gene captures | ✔ | ✔ | ✔ | ✔ | ✔ |
| Ctenotus regius | 137945 | Gene captures | ✔ | ✔ | ✔ | ✔ | ✔ |

| | | | | | | | |
|---|---|---|---|---|---|---|---|
| Ctenotus rimacolus | 30342 | Gene captures | ✔ | ✔ | ✔ | ✔ | ✔ |
| Ctenotus rubicundus | 128170 | Gene captures | ✔ | ✔ | ✔ | ✔ | ✔ |
| Ctenotus septenarius | 88059 | Gene captures | ✔ | ✔ | ✔ | ✔ | ✔ |
| Ctenotus spaldingi | 70692 | Gene captures | ✔ | ✔ | ✔ | ✔ | ✔ |

| | | | | | | | |
|---|---|---|---|---|---|---|---|
| Ctenotus strauchii | 9042 | Gene captures | ✔ | ✔ | ✔ | ✔ | ✔ |
| Ctenotus striaticeps | 30403 | Gene captures | ✔ | ✔ | ✔ | ✔ | ✔ |
| Ctenotus taeniatus | 37907 | Gene captures | ✔ | ✔ | ✔ | ✔ | ✔ |
| Ctenotus taeniolatus | 16644 | Gene captures | ✔ | ✔ | ✔ | ✔ | ✔ |
| Ctenotus vertebralis | 28180 | Gene captures | ✔ | ✔ | ✔ | ✔ | ✔ |
| Lampropholis delicata | 93599 | Gene captures | ✔ | ✔ | ✔ | ✔ | ✔ |
| Lerista aericeps | 73312 | Gene captures | ✔ | ✔ | ✔ | ✔ | ✔ |
| Lerista ameles | 77171 | Gene captures | ✔ | ✔ | ✔ | ✔ | ✔ |
| Lerista arenicola | 140003 | Gene captures | ✔ | ✔ | ✔ | ✔ | ✔ |
| Lerista axillaris | 63826 | Gene captures | ✔ | ✔ | ✔ | ✔ | ✔ |
| Lerista baynesi | 133331 | Gene captures | ✔ | ✔ | ✔ | ✔ | ✔ |
| Lerista bipes | 60733 | Gene captures | ✔ | ✔ | ✔ | ✔ | ✔ |
| Lerista borealis | 63741 | Gene captures | ✔ | ✔ | ✔ | ✔ | ✔ |
| Lerista bougainvillii | 11151 | Gene captures | ✔ | ✔ | ✔ | ✔ | ✔ |
| Lerista carpentariae | 41192 | Gene captures | ✔ | ✔ | ✔ | ✔ | ✔ |
| Lerista chalybura | 62967 | Gene captures | | | | | |
| Lerista chordae | 77000 | Gene captures | ✔ | ✔ | ✔ | ✔ | ✔ |
| Lerista cinerea | 72914 | Gene captures | ✔ | ✔ | ✔ | ✔ | ✔ |
| Lerista connivens | 59766 | Gene captures | ✔ | ✔ | ✔ | ✔ | ✔ |
| Lerista desertorum | 12580 | Gene captures | ✔ | ✔ | ✔ | ✔ | ✔ |
| Lerista dorsalis | 85818 | Gene captures | ✔ | ✔ | ✔ | ✔ | ✔ |
| Lerista edwardsae | 108252 | Gene captures | ✔ | ✔ | ✔ | ✔ | ✔ |
| Lerista elegans | 63791 | Gene captures | ✔ | ✔ | ✔ | ✔ | ✔ |
| Lerista elongata | 57634 | Gene captures | ✔ | ✔ | ✔ | ✔ | ✔ |
| Lerista emmotti | 32033 | Gene captures | ✔ | ✔ | ✔ | ✔ | ✔ |
| Lerista fragilis | 102718 | Gene captures | ✔ | ✔ | ✔ | ✔ | ✔ |
| Lerista frosti | 24169 | Gene captures | ✔ | ✔ | ✔ | ✔ | ✔ |
| Lerista gascoynensis | 138913 | Gene captures | ✔ | ✔ | ✔ | ✔ | ✔ |
| Lerista gerrardii | 63760 | Gene captures | ✔ | ✔ | ✔ | ✔ | ✔ |
| Lerista greeri | 63744 | Gene captures | ✔ | ✔ | ✔ | ✔ | ✔ |
| Lerista ips | 91444 | Gene captures | ✔ | ✔ | ✔ | ✔ | ✔ |

| | | | | | | | |
|---|---|---|---|---|---|---|---|
| Lerista karschmidti | 30712 | Gene captures | ✔ | ✔ | ✔ | ✔ | ✔ |
| Lerista kennedyensis | 63954 | Gene captures | ✔ | ✔ | ✔ | ✔ | ✔ |
| Lerista lineopunctulata | 72594 | Gene captures | ✔ | ✔ | ✔ | ✔ | ✔ |
| Lerista microtis | 63807 | Gene captures | ✔ | ✔ | ✔ | ✔ | ✔ |
| Lerista neander | 63650 | Gene captures | ✔ | ✔ | ✔ | ✔ | ✔ |
| Lerista nichollsi | 59789 | Gene captures | ✔ | ✔ | ✔ | ✔ | ✔ |
| Lerista orientalis | 29482 | Gene captures | ✔ | ✔ | ✔ | ✔ | ✔ |

| | | | | | | | | |
|---|---|---|---|---|---|---|---|---|
| Lerista picturata | 23720 | | Gene captures | ✓ | ✓ | ✓ | ✓ | ✓ |
| Lerista planiventralis | 109119 | | Gene captures | ✓ | ✓ | ✓ | ✓ | ✓ |
| Lerista praepedita | 62971 | | Gene captures | ✓ | ✓ | ✓ | ✓ | ✓ |
| Lerista terdigitata | 56833 | | Gene captures | ✓ | ✓ | ✓ | ✓ | ✓ |
| Lerista timida | 74055 | | Gene captures | | | | | |
| Lerista uniduo | 63645 | | Gene captures | ✓ | ✓ | ✓ | ✓ | ✓ |
| Lerista varia | 138923 | | Gene captures | ✓ | ✓ | ✓ | ✓ | ✓ |
| Lerista walkeri | 63823 | | Gene captures | ✓ | ✓ | ✓ | ✓ | ✓ |
| Lerista wilkinsi | 76998 | | Gene captures | ✓ | ✓ | ✓ | ✓ | ✓ |
| Anolis carolinensis | | XP_008102123.1, AF134189/90/91, AH007735.2, AH007736.2, AF133907.1 | NCBI | ✓ | ✓ | ✓ | ✓ | ✓ |
| Bos taurus | | NM_174566, NM_0010148, NM_174567 | NCBI | ✓ | ✓ | | ✓ | |
| Carlia fusca | | AY508937 | NCBI | | | ✓ | | |
| Carlia rhomboidalis | | AY508933 | NCBI | | | ✓ | | |
| Carlia rostralis | | AY508939 | NCBI | | | ✓ | | |
| Carlia rubrigularis | | AY508932 | NCBI | | | ✓ | | |
| Carlia rufilatus | | AY508938 | NCBI | | | ✓ | | |
| Carlia timlowi | | AY508935 | NCBI | | | ✓ | | |
| Carlia vivax | | AY508934 | NCBI | | | ✓ | | |
| Feylinia sp | | KR336714, KR336742, KR336754, KR336717, KR336751 | NCBI | ✓ | ✓ | ✓ | ✓ | ✓ |
| Lampropholis coggeri | | AY508941 | NCBI | | | ✓ | | |
| Melanoseps occidentalis | | KR336713.1, KR336743.1, KR336718.1, KR336750 | NCBI | ✓ | ✓ | | ✓ | ✓ |
| Python bivittatus | | LOC103048730, XP_007423324.1, XP_007441698.1 | NCBI | ✓ | ✓ | | ✓ | |
| Saproscincus basiliscus | | AY508940 | NCBI | | | ✓ | | |
| Sphenodon punctatus | | | NCBI | ✓ | ✓ | ✓ | ✓ | ✓ |
| Anomalopus brevicollis | | | transcriptomes | ✓ | | | | |
| Chalcides ocellatus | | | transcriptomes | ✓ | ✓ | ✓ | ✓ | |
| Ctenotus atlas | | | transcriptomes | ✓ | ✓ | ✓ | ✓ | ✓ |
| Eumeces schneideri | | | transcriptomes | ✓ | ✓ | ✓ | ✓ | ✓ |
| Eutropis macularia | | | transcriptomes | ✓ | | ✓ | ✓ | |
| Glaphyromorphus punctulatus | | | transcriptomes | ✓ | ✓ | ✓ | ✓ | ✓ |
| Lerista dorsalis | | | transcriptomes | ✓ | ✓ | ✓ | ✓ | ✓ |
| Lerista edwardsae | | | transcriptomes | ✓ | ✓ | ✓ | ✓ | ✓ |
| Lerista terdigitata | | | transcriptomes | ✓ | ✓ | ✓ | ✓ | ✓ |
| Liopholis inornata | | | transcriptomes | ✓ | ✓ | ✓ | ✓ | ✓ |
| Melanoseps occidentalis | | | transcriptomes | ✓ | | | | ✓ |
| Platysaurus broadleyi | | | transcriptomes | ✓ | ✓ | ✓ | ✓ | ✓ |
| Tiliqua rugosa | | | transcriptomes | ✓ | | | | |

### Table 2. Site CodeML results

| Models | Parameters | D.F. | Models Compared | 2Δ (ln L) | P | |
|---|---|---|---|---|---|---|
| **1. *lws* opsin gene** | | | | | | |
| A. M1a | ω0=0.031, ω1=1, p0=0.924, p1=0.076 | | | | | |
| B. M2a | ω0=0.035, ω1=1, ω2=3.753, p0=0.925, p1=0.055, p2=0.020 | 2 | B vs. A | 57.398 | 0 | ** |
| C. M7 | p=0.025, q=0.133 | | | | | |
| D. M8 (β&ω) | p0=0.977, p1=0.023, p=0.103, q=1.007, ω=3.482 | 2 | D vs. C | 64.489 | 0 | ** |
| **2. *sws1* opsin gene** | | | | | | |
| E. M1a | ω0=0.022, ω1=1, p0=0.948, p1=0.052 | | | | | |
| F. M2a | ω0=0.022, ω1=1, ω2=15.951, p0=0.948, p1=0.052, p2=0.000 | 2 | F vs. E | 2.60E-05 | 1 | |
| G. M7 | p=0.084, q=0.854 | | | | | |
| H. M8 (β&ω) | p0=0.955, p1=0.045, p=0.334, q=11.794, ω=1.000 | 2 | H vs G | 22.05 | 0.000016 | ** |
| **3. *rh1* rhodopsin gene** | | | | | | |
| I. M1a | ω0=0.024, ω1=1, p0=0.848, p1=0.152 | | | | | |
| J. M2a | ω0=0.024, ω1=1, ω2=1, p0=0.849, p1=0.102, p2=0.049 | 2 | J vs. I | 0.881 | 0.644 | |
| K. M7β | p=0.071, q=0.387 | | | | | |
| L. M8 (β&ω) | p0=0.945, p1=0.055, p=0.105, q=0.1.033, ω=1.436 | 2 | L vs. K | 5.017 | 0.081 | |
| **3. *rh2* rhodopsin gene** | | | | | | |
| M. M1a | ω0=0.015, ω1=1, p0=0.855, p1=0.145 | | | | | |
| N. M2a | ω0=0.016, ω1=1, ω2=1.587, p0=0.861, p1=0.104, p2=0.035 | 2 | N vs. M | 7.432 | 0.024 | * |
| O. M7β | p=0.019, q=0.097 | | | | | |
| P. M8 (β&ω) | p0=0.980, p1=0.020, p=0.028, q=0.160, ω=1.828 | 2 | P vs. O | 8.128 | 0.017 | * |
| **3. *sws2* rhodopsin gene** | | | | | | |
| Q. M1a | ω0=0.026, ω1=1, p0=0.812, p1=0.188 | | | | | |
| R. M2a | ω0=0.029, ω1=1, ω2=2.733, p0=0.816, p1=0.143, p2=0.040 | 2 | R vs. Q | 27.862 | 0.000001 | ** |
| S. M7β | p=0.027, q=0.139 | | | | | |
| T. M8 (β&ω) | p0=0.950, p1=0.050, p=0.085, q=0.520, ω=2.482 | 2 | T vs. S | 36.696 | 0 | ** |

#### Table 3. Branch CodeML Results

| Models | $\omega(d_N/d_S)$ | D.F. | Models Compared | 2Δ (in *L*) | P |
|---|---|---|---|---|---|
| **1. *sws1* opsin gene** | | | | | |
| A. All branches have one ω | ω=0.0475 | - | - | - | - |
| B. Temperate, ω1; Arid, ω2 | ω1=0.0582, ω2=0.0402 | 1 | B vs. A | 0.291 | 0.589 |
| C. Mobile eyelids, ω1; Fused Lower Eyelid, ω2 | ω1=0.0001, ω2=0.0565 | 1 | C vs. A | 1.104 | 0.293 |
| D. <3 Hindlimb Digits, ω1; ≥3 Hindlimb Digits, ω2 | ω1=0.0561, ω2=0.0201 | 1 | D vs. A | 0.969 | 1 |
| E. Each branch has its own ω | Variable by branch | 38 | E vs. A | 30.73 | 0.999 |
| **2. *lws* opsin gene** | | | | | |
| F. All branches have one ω | ω=0.120 | - | - | - | - |
| G. Temperate, ω1; Arid, ω2 | ω1=0.110, ω2=0.132 | 1 | G vs. F | 0.269 | 0.604 |
| H. Mobile eyelids, ω1; Fused Lower Eyelid, ω2 | ω1=0.0723, ω2=0.140 | 1 | H vs. F | 2.49 | 0.115 |
| I. <3 Hindlimb Digits, ω1; ≥3 Hindlimb Digits, ω2 | ω1=0.151, ω2=0.0664 | 1 | I vs. F | 4.249 | 0.0393 |
| J. Each branch has its own ω | Variable by branch | 38 | J vs. F | 46.412 | 0.996 |
| **3. *rh1* rhodopsin gene** | | | | | |
| K. All branches have one ω | ω=0.125 | - | - | - | - |
| L. Temperate, ω1; Arid, ω2 | ω1=0.147, ω2=0.104 | 1 | L vs. K | 2.703 | 0.1 |
| M. Mobile eyelids, ω1; Fused Lower Eyelid, ω2 | ω1=0.124, ω2=0.125 | 1 | M vs. K | 0.00231 | 0.962 |
| N. <3 Hindlimb Digits, ω1; ≥3 Hindlimb Digits, ω2 | ω1=0.119, ω2=0.139 | 1 | N vs. K | 0.48 | 0.489 |
| O. Each branch has its own ω | Variable by branch | 38 | O vs. K | 23.935 | 1 |
| **3. *rh2* rhodopsin gene** | | | | | |
| P. All branches have one ω | ω=0.464 | - | - | - | - |
| Q. Temperate, ω1; Arid, ω2 | ω1=0.668, ω2=0.344 | 1 | Q vs. P | 2.1 | 0.147 |
| R. Mobile eyelids, ω1; Fused Lower Eyelid, ω2 | ω1=0.360, ω2=0.482 | 1 | R vs. P | 0.182 | 0.67 |
| S. <3 Hindlimb Digits, ω1; ≥3 Hindlimb Digits, ω2 | ω1=0.437, ω2=0.524 | 1 | S vs. P | 0.143 | 0.705 |
| T. Each branch has its own ω | Variable by branch | 38 | T vs. P | 38.444 | 0.999 |
| **3. *sws2* rhodopsin gene** | | | | | |
| U. All branches have one ω | ω=0.260 | - | - | - | - |
| V. Temperate, ω1; Arid, ω2 | ω1=0.258, ω2=0.262 | 1 | V vs. U | 0.00293 | 0.957 |
| W. Mobile eyelids, ω1; Fused Lower Eyelid, ω2 | ω1=0.234, ω2=0.267 | 1 | W vs. U | 0.15 | 0.698 |
| X. <3 Hindlimb Digits, ω1; ≥3 Hindlimb Digits, ω2 | ω1=0.248, ω2=0.288 | 1 | X vs. U | 0.273 | 0.601 |
| Y. Each branch has its own ω | Variable by branch | 38 | Y vs. U | 47.696 | 0.994 |

#### Table 4. Branch-site CodeML Results

| Gene | Foreground branch | Sites under positive selection | 2Δl | P Value |
|---|---|---|---|---|
| ***lws* opsin gene** | Arid | 112 | 10.91 | **0.000956** |
| | Fixed Lower Eyelid | None | 0.000066 | **0.994** |
| | <3 Hindlimb Digits | 112- 270 | 47.853 | 0 |
| ***sws1* opsin gene** | Arid | None | 2.07 | **1** |
| | Fixed Lower Eyelid | None | 0.465 | **1** |
| | <3 Hindlimb Digits | None | 0.229 | 1 |
| ***rh1* rhodopsin gene** | Arid | None | 0.0448 | 0.832 |
| | Fixed Lower Eyelid | None | 0.0145 | **0.904** |
| | <3 Hindlimb Digits | None | 0.0195 | 1 |
| ***rh2* rhodopsin gene** | Arid | 72 - 79 - 96 - 113 | 29.073 | 0 |
| | Fixed Lower Eyelid | None | 0 | **1** |
| | <3 Hindlimb Digits | 72 - 79 - 96 | 44.954 | 0 |
| ***sws2* opsin gene** | Arid | 2 - 108 | 16.532 | 0.000048 |
| | Fixed Lower Eyelid | None | 0 | **1** |
| | <3 Hindlimb Digits | 2 - 11 - 50 - 88 - 89 - 108 | 57.734 | 0 |

### Table 5. Clade CodeML Results

| *lws* opsin genes | | | | | | | | | | | | |
|---|---|---|---|---|---|---|---|---|---|---|---|---|
| **Partition** | **Model** | **NP** | **ln*L*** | **AIC** | ***k*** | **Parameters** | | | **Null** | **2Δl** | **df** | **P** |
| | | | | | | **ω0** | **ω1** | **ω2** | | | | |
| — | M2a_rel | 81 | -2534.343169 | 5230.686338 | 1.82447 | 0.00466 (0.95942) | 1 (0.03240) | 15.24374 (0.00818) | — | — | — | — |
| Arid | CmC | 82 | -2569.302 | 5302.604 | 1.70265 | 0 (0) | 1 (0.03931) | 0.00876 (0.96069) | M2a_rel | 69.91766 | 1 | **6.18E-17** |
| Tropical/ Temperate | CmC | 82 | -2569.302 | 5302.604 | 1.70265 | 0 (0) | 1 (0.03931) | 0 (0.96069) | M2a_rel | 69.91766 | 1 | **6.18E-17** |
| Fused Eyelids | CmC | 82 | -2569.327422 | 5302.654844 | 1.69896 | 0 (0.0001) | 1 (0.0381) | 0 (0.9619) | M2a_rel | 69.96851 | 1 | **6.03E-17** |
| Unfused Eyelids | CmC | 82 | -2569.327422 | 5302.654844 | 1.69896 | 0 (0.0001) | 1 (0.0381) | 0.00732 (0.9619) | M2a_rel | 69.96851 | 1 | **6.03E-17** |
| <3 Hindlimb Digits | CmC | 82 | -2569.126441 | 5302.252882 | 1.69608 | 0 (0) | 1 (0.03649) | 0.00921 (0.96351) | M2a_rel | 69.56654 | 1 | **7.39E-17** |
| ≥3 Hindlimb Digits | CmC | 82 | -2569.126441 | 5302.252882 | 1.69608 | 0 (0) | 1 (0.03649) | 0 (0.96351) | M2a_rel | 69.56654 | 1 | **7.39E-17** |

| *rh1* rhodopsin gene | | | | | | | | | | | | |
|---|---|---|---|---|---|---|---|---|---|---|---|---|
| **Partition** | **Model** | **NP** | **lnL** | **AIC** | **k** | **Parameters** | | | **Null** | **2Δl** | **df** | **P** |
| | | | | | | **ω0** | **ω1** | **ω2** | | | | |
| — | M2a_rel | 81 | -2827.813523 | 5817.627046 | 3.91093 | 0.00598 (0.90112) | 1 (0.06925) | 2.32614 (0.02963) | — | — | — | — |
| Arid | CmC | 82 | -2831.171943 | 5826.343886 | 3.64271 | 0.00413 (0.89120) | 1 (0.073) | 0.77616 (0.0358) | M2a_rel | 6.71684 | 1 | **9.55E-03** |
| Tropical/ Temperate | CmC | 82 | -2831.171943 | 5826.343886 | 3.64271 | 0.00413 (0.89120) | 1 (0.073) | 0.10366 (0.03580) | M2a_rel | 6.71684 | 1 | **9.55E-03** |
| Fused Eyelids | CmC | 82 | -2837.809655 | 5839.61931 | 3.91063 | 0.00598 (0.90113) | 1 (0.06905) | 2.40908 (0.02981) | M2a_rel | 19.99226 | 1 | **7.78E-06** |
| Unfused Eyelids | CmC | 82 | -2837.809655 | 5839.61931 | 3.91063 | 0.00598 (0.90113) | 1 (0.06905) | 2.30557 (0.02981) | M2a_rel | 19.99226 | 1 | **7.78E-06** |
| <3 Hindlimb Digits | CmC | 82 | -2833.762294 | 5831.524588 | 3.70533 | 0.03671 (0.11346) | 1 (0.09715) | 0 (0.78938) | M2a_rel | 11.89754 | 1 | **5.62E-04** |
| ≥3 Hindlimb Digits | CmC | 82 | -2833.762294 | 5831.524588 | 3.70533 | 0.03671 (0.11346) | 1 (0.09715) | 0.00602 (0.78938) | M2a_rel | 11.89754 | 1 | **5.62E-04** |

***rh2* opsin gene**

| Partition | Model | NP | lnL | AIC | k | Parameters | | | Null | 2Δl | df | P |
|---|---|---|---|---|---|---|---|---|---|---|---|---|
| | | | | | | **ω0** | **ω1** | **ω2** | | | | |
| — | M2a_rel | 81 | -1095.233723 | 2352.467446 | 2.32473 | 0.03085 (0.93225) | 1 (0.04225) | 15.40160 (0.0255) | — | — | — | — |
| Arid | CmC | 82 | -1139.549363 | 2443.098726 | 1.77554 | 0 (0) | 1 (0.0808) | 0.01804 (0.9192) | M2a_rel | 88.63128 | 1 | **4.76E-21** |
| Tropical/ Temperate | CmC | 82 | -1139.549363 | 2443.098726 | 1.77554 | 0 (0) | 1 (0.0808) | 0 (0.9192) | M2a_rel | 88.63128 | 1 | **4.76E-21** |
| Fused Eyelids | CmC | 82 | -1139.784896 | 2443.569792 | 1.76989 | 0 (0) | 1 (0.08132) | 0 (0.91868) | M2a_rel | 89.10235 | 1 | **3.75E-21** |
| Unfused Eyelids | CmC | 82 | -1139.784896 | 2443.569792 | 1.76989 | 0 (0) | 1 (0.08132) | 0.00909 (0.91868) | M2a_rel | 89.10235 | 1 | **3.75E-21** |
| <3 Hindlimb Digits | CmC | 82 | -1138.274821 | 2440.549642 | 1.81701 | 0.00538 (0.91632) | 1 (0.0762) | 0 (0.00748) | M2a_rel | 86.0822 | 1 | **1.73E-20** |
| ≥3 Hindlimb Digits | CmC | 82 | -1138.274821 | 2440.549642 | 1.81701 | 0.00538 (0.91632) | 1 (0.0762) | 8.05017 (0.00748) | M2a_rel | 86.0822 | 1 | **1.73E-20** |

***sws1* opsin gene**

| Partition | Model | NP | lnL | AIC | k | Parameters | | | Null | 2Δl | df | P |
|---|---|---|---|---|---|---|---|---|---|---|---|---|
| | | | | | | **ω0** | **ω1** | **ω2** | | | | |
| — | M2a_rel | 81 | -1835.272374 | 3832.544748 | 9.1945 | 0.00005 (0.98586) | 1 (0.00110) | 4.45544 (0.01304) | — | — | — | — |
| Arid | CmC | 82 | -1838.917434 | 3841.834868 | 8.7285 | 0 (0.8719) | 1 (0.02554) | 0 (0.10256) | M2a_rel | 7.29012 | 1 | **6.93E-03** |
| Tropical/ Temperate | CmC | 82 | -1838.917434 | 3841.834868 | 8.7285 | 0 (0.8719) | 1 (0.02554) | 0 (0.10256) | M2a_rel | 7.29012 | 1 | **6.93E-03** |
| Fused Eyelids | CmC | 82 | -1840.283982 | 3844.567964 | 8.72495 | 0 (0.87727) | 1 (0.02555) | 0 (.09719) | M2a_rel | 10.02322 | 1 | **1.55E-03** |
| Unfused Eyelids | CmC | 82 | -1840.283982 | 3844.567964 | 8.72495 | 0 (0.87727) | 1 (0.02555) | 0 (.09719) | M2a_rel | 10.02322 | 1 | **1.55E-03** |
| <3 Hindlimb Digits | CmC | 82 | -1838.917434 | 3841.834868 | 8.72848 | 0 (0.83713) | 1 (0.02554) | 0 (0.13733) | M2a_rel | 7.29012 | 1 | **6.93E-03** |
| ≥3 Hindlimb Digits | CmC | 82 | -1838.917434 | 3841.834868 | 8.72848 | 0 (0.83713) | 1 (0.02554) | 0 (0.13733) | M2a_rel | 7.29012 | 1 | **6.93E-03** |

*sws2* opsin gene

| Partition | Model | NP | lnL | AIC | k | Parameters | | | Null | 2Δl | df | p |
|---|---|---|---|---|---|---|---|---|---|---|---|---|
| | | | | | | ω0 | ω1 | ω2 | | | | |
| — | M2a_rel | 81 | -1723.931778 | 3609.863556 | 7.37316 | 0.01186 (0.91176) | 1 (0.04098) | 6.58461 (0.04727) | — | — | — | — |
| Arid | CmC | 82 | -1779.143238 | 3722.286476 | 5.43399 | 0 (0.85413) | 1 (0.08704) | 0.5061 (0.05883) | M2a_rel | 110.4229 | 1 | **7.92E-26** |
| Tropical/ Temperate | CmC | 82 | -1779.143238 | 3722.286476 | 5.43399 | 0 (0.85413) | 1 (0.08704) | 0 (0.05883) | M2a_rel | 110.4229 | 1 | **7.92E-26** |
| Fused Eyelids | CmC | 82 | -1780.517123 | 3725.034246 | 5.39046 | 0 (0) | 1 (0.08803) | 0 (0.91197) | M2a_rel | 113.1707 | 1 | **1.98E-26** |
| Unfused Eyelids | CmC | 82 | -1780.517123 | 3725.034246 | 5.39046 | 0 (0) | 1 (0.08803) | 0.00992 (0.91197) | M2a_rel | 113.1707 | 1 | **1.98E-26** |
| <3 Hindlimb Digits | CmC | 82 | -1781.311377 | 3726.622754 | 5.41024 | 0.00579 (0.00001) | 1 (0.09) | 0.00834 (0.90999) | M2a_rel | 114.7592 | 1 | **8.89E-27** |
| ≥3 Hindlimb Digits | CmC | 82 | -1781.311377 | 3726.622754 | 5.41024 | 0.00579 (0.00001) | 1 (0.09) | 0.00555 (0.90999) | M2a_rel | 114.7592 | 1 | **8.89E-27** |

### Table 6. Genes Sampled with Probes and Probe Used

| Gene symbol | Description | Function | Probe Sequence Species | Number of Probe Species | BC/BS | Size (bp) in *Anolis* |
|---|---|---|---|---|---|---|
| LWS (OPN1LW) | Long wavelength sensitive opsin | Visual Opsin | *Anolis, Chrysemys, Gallus, Gecko, Python, Thamnophis* | 6 | BC/BS | 1110 |
| RH1 (RHO) | Rhodopsin 1 | Visual Opsin | *Alligator, Anolis, Gallus, Pelodiscus, Thamnophis, Uta* | 6 | BC/BS | 1059 |
| RH2 | Rhodopsin 2 | Visual Opsin | *Anolis, Chrysemys, Gallus, Gekko, Pelodiscus, Uta* | 6 | BC/BS | 1068 |
| SWS1 (OPN1SW) | Short-wave sensitive opsin 1 | Visual Opsin | *Anolis, Chrysemys, Gallus, Gecko, Python, Thamnophis, Uta* | 7 | BC/BS | 1041 |
| SWS2 | Short-wave sensitive opsin 2 | Visual Opsin | *Alligator, Anolis, Chrysemys, Gallus, Uta* | 5 | BC/BS | 1092 |

#### Table 7. RELAX Results

| | Climate | Eyelids | <3 hindlimb digits |
|---|---|---|---|
| LWS | p=0.2664<br>k=1.39 | p=0.2628<br>k=1.34 | p=0.0141<br>k=0.73 |
| RH1 | p=0.0910<br>k=0.84 | p=0.9542<br>k=1.01 | p=0.5001<br>k=1.08 |
| RH2 | p=0.1477<br>k=0.32 | p=0.8418<br>k=9.45 | p=0.7091<br>k=1.29 |
| SWS1 | p=0.5892<br>k=0.82 | p=0.0742<br>k=12.03 | p=0.2708<br>k=0.18 |
| SWS2 | p=0.9915<br>k=1.02 | p=0.8831<br>k=0.95 | p=0.4751<br>k=2.02 |
